# Loss of starch synthase IIa alleviates the negative impact of high temperature on rice starch during grain filling

**DOI:** 10.64898/2026.09.28.754565

**Authors:** Shumpei Hashimoto, Ryo Matsushima, Naoko F. Oitome, Yuko Hosaka, Satoko Miura, Naoko Crofts, Toshio Yamamoto, Naoko Fujita

## Abstract

High temperatures during grain filling stage are becoming increasingly frequent, compromising both grain and eating quality and thereby driving demand for heat-resilient cultivars. Such conditions are known to reduce the expression of *granule-bound starch synthase I* (*GBSSI*) and *starch branching enzyme IIb* (*BEIIb*), which are involved in starch biosynthesis, resulting in a decrease in amylose content and an increase in long-chain amylopectin. Thus, the present study introduced functional mutation in *starch synthase IIa* (*SSIIa*) that increases the proportion of short amylopectin chains to genetically compensate for the high-temperature-induced increase in amylopectin long chain. Rice lines carrying the *ss2a* mutation were grown at two locations with cooler (Akita) and warmer (Okayama) temperatures. Their grain traits, starch structure, and eating quality were compared. The *ss2a* mutant lines showed an increased proportion of short amylopectin chains as well as an increased apparent amylose content. Furthermore, these alterations in starch structure varied with the grain-filling temperature of the cultivation sites, ultimately affected eating quality. These results suggest that enriching short amylopectin chain via the *ss2a* mutation can counteract the increase in long amylopectin chain caused by high temperatures during grain filling, thereby maintaining a desirable starch structure and eating quality.

## Introduction

In recent years, high temperatures associated with global warming have seriously impacted agricultural production and food security worldwide (Gitz and Meybeck 2016, Roberts and Schlenker 2013, Shukla *et al*. 2019). Rice (*Oryza sativa* L.) is one of the world’s most important cereal crops and serves as a staple food for more than half of the global population. Ensuring stable rice production while maintaining grain quality is therefore a critical challenge for global food security (Li *et al*. 2023). In temperate regions, including Japan, high temperatures during grain filling stage have led grain quality deterioration due to heat stress (Itoh *et al*. 2024). Consequently, there is an urgent need to develop rice cultivars that can stably maintain not only grain yield but also desirable appearance and eating quality under high-temperature conditions.

Rice endosperm starch consists of amylose, composed primarily of linear glucan chains, and amylopectin, a highly branched glucan polymer. Amylopectin accounts for approximately 80% of rice starch, and its biosynthesis involves multiple enzymes and their isozymes that coordinately regulate glucan chain elongation, branching, and debranching(Nakamura 2002). Specifically, starch synthases (SSs) elongate linear glucan chains through α-1,4 glycosidic linkages using ADP-glucose as a substrate; starch branching enzymes (BEs) cleave linear glucan chains and introduce new branches through α-1,6 glycosidic linkages; and starch debranching enzymes, including isoamylase and pullulanase, remove inappropriate branches(Nakamura 2002). Each enzyme class comprises multiple isozymes that differ in tissue specificity(Koller *et al*. 2002, Yamanouchi and Nakamura 1992), temporal expression patterns(Ohdan *et al*. 2005), and substrate preferences for glucan chains(Fujita *et al*. 2011, Hwang *et al*. 2010, Nakamura *et al*. 2010). In the developing rice endosperm, three SS isozymes (SSI, SSIIa and SSIIIa) and two BE isozymes (BEI and BEIIb) play major roles in amylopectin biosynthesis. Short branches with a degree of polymerization (DP) of 6–7 generated by BEIIb(Nakamura *et al*. 2014) are elongated to DP 8–12 by SSI(Fujita *et al*. 2006). SSIIa further elongates short amylopectin chains with DP ≤ 12 to intermediate chains of 13 ≤ DP ≤ 24(Nakamura *et al*. 2005), whereas SSIIIa is primarily involved in the synthesis of longer amylopectin chains with DP > 30 connecting amylopectin clusters(Fujita *et al*. 2007). These enzymes do not function independently. For example, SSIIa forms a trimeric complex with SSI and BEIIb in maize(Hennen-Bierwagen *et al*. 2008, Liu *et al*. 2009) and rice (Crofts *et al*. 2015, Ying *et al*. 2022), and this complex is thought to play an important role in the biosynthesis of short and intermediate amylopectin chains (DP ≤ 24) within amylopectin clusters.

One of the most prominent forms of grain quality deterioration caused by high temperatures during grain filling is an increased occurrence of chalky grains(Morita *et al*. 2016, Wada *et al*. 2019). In chalky grains, insufficient packing of starch granules in the endosperm creates microscopic air spaces between the granules, resulting in diffuse light reflection and an opaque appearance in part or all of the grain(Nevame *et al*. 2018, Tashiro and Wardlaw 1991). Under high-temperature conditions, the balance between starch accumulation and degradation in the endosperm is altered, and increased expression of starch-degrading enzymes, particularly α-amylases, has been shown to contribute to the formation of chalky grains(Hakata *et al*. 2012, Nakata *et al*. 2017). Enhancing substrate supply for starch biosynthesis may therefore represent an effective breeding strategy for preventing the reduction in starch accumulation under high-temperature conditions. Introduction of the indica rice *sucrose synthase 3* (*Sus3*) allele into japonica rice cultivars has been reported to maintain carbon supply for starch biosynthesis during grain filling under high-temperature conditions and substantially reduce the occurrence of chalky grains(Murata *et al*. 2014, Takehara *et al*. 2018). Heat-tolerant cultivars such as ‘Fu-Fu-Fu’ have subsequently been developed by introducing *Sus3* from indica rice, demonstrating that maintaining sufficient starch accumulation is an effective strategy for mitigating the deterioration of grain appearance caused by high temperatures during grain filling(Goto *et al*. 2026, Murata *et al*. 2022).

However, the adverse effects of high temperatures during grain filling are not limited to the deterioration of grain appearance represented by chalkiness. Grain-filling temperature also has a profound effect on starch structure. When the japonica rice cultivar ‘Akitakomachi’ was subjected to a high grain-filling temperature with a mean temperature of 28°C, both amylose content and the proportion of short amylopectin chains were significantly reduced compared with those under a control temperature of 22°C, at which most grains developed normally(Kato *et al*. 2019). These changes are thought to be associated with reduced expression of *granule-bound starch synthase I* (*GBSSI*), which is primarily responsible for amylose biosynthesis, and *BEIIb*, which plays a major role in determining the branched structure of amylopectin, respectively(Kato *et al*. 2019, Larkin and Park 1999). A moderate reduction in amylose content can generally soften cooked rice and, in some cases, improve its eating quality. However, excessive reductions in amylose content and short amylopectin chains can alter starch gelatinization and physical properties of cooked rice. In particular, the shift toward longer amylopectin chains caused by reduced *BEIIb* expression contributes to the deterioration of eating quality under high-temperature conditions(Kato *et al*. 2019).

Importantly, the mechanisms underlying heat-induced chalky grain formation and changes in starch molecular structure are fundamentally distinct. Thus, even when chalky grain formation is successfully suppressed, the reduction in short amylopectin chains caused by high-temperature grain filling may persist, and improvement in eating quality cannot necessarily be expected. Comprehensive improvement of rice quality under high-temperature grain-filling conditions therefore requires not only the suppression of chalky grain formation but also a new breeding strategy for maintaining a desirable starch structure under heat stress.

The target of the present study is SSIIa, one of key enzymes regulating amylopectin chain-length distribution. The SSIIa haplotype commonly found in japonica rice cultivars carries three single-nucleotide polymorphisms that substantially reduce SSIIa activity, resulting in a higher proportion of short amylopectin chains with DP ≤ 12 than that in indica rice cultivars(Nakamura *et al*. 2005, Umemoto *et al*. 2002). More recently, a complete loss-of-function mutant of *SSIIa* was identified, and loss of SSIIa function was shown to further increase both the proportion of short amylopectin chains and apparent amylose content beyond the levels observed in conventional japonica cultivars(Miura *et al*. 2018). Notably, these alterations are opposite in direction to the reduction in short amylopectin chains and amylose content induced by high temperatures during grain filling. Therefore, it was hypothesized that loss of SSIIa could shift starch structure in a direction opposite to that induced by high temperature, thereby compensating for heat-induced changes in amylopectin structure and helping to maintain desirable starch properties and eating quality. To test this hypothesis, two rice lines carrying a loss-of-function mutation in *SSIIa* were grown at two locations with distinct temperatures during the grain-filling period. Grain appearance and seed traits, amylose content, amylopectin chain-length distribution, and eating quality were then compared to determine how the *ss2a* mutation affects starch structure and rice grain quality under different grain-filling temperatures. This enabled the evaluation of the potential of genetic modification of starch structure as a strategy for adaptation to high-temperature grain filling.

## Materials and Methods

### Plant materials and growth condition

Two backcrossed inbred lines (BILs), K19 and A21, carrying the *ss2a* allele, were used in this study.

These lines were developed by introducing the *ss2a* allele from the ‘Kinmaze’ mutant EM204 (Miura et al. 2018) into elite-eating-quality rice cultivar ‘Akitakomachi’ and high yield-large grain ‘Akita 63’, respectively (BC3F7). Through three successive backcrosses with the respective recurrent parents, the *ss2a* allele was selected by dCAPs marker with the following primers: forward, 5′-CAGACAGGTGAAGCTTCTATCTG-3′; reverse, 5′-CAAAACAGAATCATGCGCTTCATGGGTTC-3′. The amplified PCR products were digested with NlaIV. Homozygous plants for the *ss2a* allele were selected to establish K19 and A21. BIL lines were grown in 2024 at two field locations: Katagami City, Akita Prefecture, Japan (39°51′N, 140°00′E), and Kurashiki City, Okayama Prefecture, Japan (34°35′N, 133°46′E). ‘Akitakomachi’ and ‘Akita 63’ were also grown in Akita.

### Purification of starch granules from rice endosperm

Starch granules were purified from mature rice endosperm. Dehulled rice grains were polished to 80– 90% of their original weight. Three grams of the polished rice grains were soaked in 50 mL of 0.1% (w/v) NaOH overnight at 4°C. After removal of the supernatant, the grains were ground using a pre-chilled mortar and pestle on ice. The homogenate was filtered through a 100-μm nylon mesh while adding 0.1% (w/v) NaOH. The filtrate was adjusted to a final volume of 45 mL with 0.1% (w/v) NaOH and shaken on ice for 3 h, followed by incubation overnight at 4°C without agitation. The suspension was neutralized by adding 1 mL of 1 N HCl and then centrifuged at 3,000 rpm at 4°C. After removal of the supernatant, the pellet was resuspended in 40 mL of distilled water and centrifuged again under the same conditions. This washing procedure was repeated three times. The resulting starch pellet was dried under reduced pressure and stored at −30°C until further analysis.

### Chain-length distribution of amylopectin

Purified starch (3 mg) was suspended in 300 μl of 0.25 M NaOH. The suspension was boiled for 5 min. The gelatinized polyglucan sample was added by 9.6 μl of 100% acetic acid, 100 μl of 600 mM Na-acetate buffer (pH 4.4), 15 μl of 2% NaN3 and 1090 μl of distilled water. The sample was debranched by adding 6 μl of *Pseudomonas amyloderamosa* isoamylase (354 units, Nagase Viita, Okayama, Japan) at 37°C for 24 h. The hydrolyzed sample was boiled for 20 min and centrifuged. The supernatant was deionized by filtration of ion exchange resin (Bio-Rad AG 501-X8(D)) in microtube. Fluorescence labeling and capillary electrophoresis were performed according to the method of O’Shea and Morell(O’Shea and Morell 1996) and the protocols provided from a manufacturer by using the eCAP N-linked oligosaccharide profiling kit and capillary electrophoresis (P/ACE MDQ Carbohydrate System, AB Sciex).

### Apparent amylose content and the ratio of amylopectin short and long chains

Apparent amylose content and the ratio of amylopectin short and long chains were determined by gel filtration chromatography (Toyopearl HW-55S and HW-50S × 3) according to previously published methods(Fujita *et al*. 2007, Toyosawa *et al*. 2016). Gel filtration chromatography of debranched starch revealed three peaks (Fractions I, II, and III; Supplementary Fig. 1). The proportion of Fr. I was defined as apparent amylose content, and Fr. III/II ratio was used as an index of the proportion of short amylopectin chains.

### Thermal properties of starch

The thermal properties of purified starch were analyzed by DSC (LabSolutions TA software version 1.01, DSC 60 Plus; Shimadzu, Kyoto, Japan). Briefly, 3 mg of purified starch was suspended in 9 μL of distilled water and encapsulated in an aluminum seal pan (S201-53090; Shimadzu, Kyoto, Japan). Then, 45 mg of activated alumina was encapsulated in the aluminum seal pan and used as a reference sample. The thermal properties of starch were analyzed using the temperature program described previously (Fujita *et al*. 2006, 2009). The heating rate was 5°C min^-1^ over a temperature range of 5°C to 90°C.

### Sensory evaluation of cooked rice

Sensory evaluation of cooked rice was conducted as a single-blind test by 12 panelists (six females and six males; 18–45 years of age). Mature rice grains were dehulled and polished to approximately 90% of their original weight using a rice mill (Magic Mill RSKM5D, Satake, Hiroshima, Japan). The polished rice was cooked with 1.2 volumes of water in an electric rice cooker (JPF-G055, Tiger Corporation, Osaka, Japan). Cooked rice samples were evaluated for appearance, aroma, hardness, mouthfeel, stickiness and taste. Each attribute was scored on a 5-point scale, with ‘Akitakomachi’ used as the reference cultivar and assigned a score of 3. Water was provided to the panelists for palate cleansing between samples. For the evaluation of freshly cooked rice, samples were evaluated immediately after cooking. For cooled rice, freshly cooked rice was divided into small portions and cooled at approximately 15°C for 2 h before sensory evaluation. The sensory score for each attribute was calculated as the mean score obtained from the 12 panelists.

## Results

### Plant materials and growth location for evaluating high-temperature grain filling

To evaluate the effects of the *ss2a* mutation on grain quality and starch properties under high-temperature grain-filling conditions, two backcrossed inbred lines (BILs) with different genetic backgrounds were used (Fig. 1A). ‘Akitakomachi’, one of the background cultivars, is a widely cultivated Japanese rice cultivar known for its high eating quality. In contrast, ‘Akita 63’ is a high-yielding rice cultivar characterized by large grains. The two BILs, K19 and A21, were developed by introducing a loss-of-function *ss2a* allele derived from an *ss2a* mutant in the ‘Kinmaze’ genetic background through three rounds of backcrossing. In the loss-of-function allele, the last nucleotide of intron 5 of the *SSIIa* gene is substituted from guanine (G) to adenine (A) (Fig. 1B). This single-nucleotide substitution results in the removal of exon 6 together with introns 5 and 6 during splicing, generating a 45-bp deletion in the mature mRNA. Consequently, the SSIIa protein encoded by this allele lacks 15 amino acids (Miura *et al*. 2018). These lines were grown in 2024 at two contrasting locations in Katagami City, Akita Prefecture (39°43′N, 140°06′E), and Kurashiki City, Okayama Prefecture (34°35′N, 133°46′E) (Fig. 1C, D). Because of their different geographical locations, the two sites differed in temperature during the growing season, with Okayama generally experiencing higher temperatures than Akita (Fig. 1E). During the grain-filling period from August to early September, temperatures in Okayama were approximately 2–4°C higher than those in Akita (Fig. 1F).

**Fig 1.**
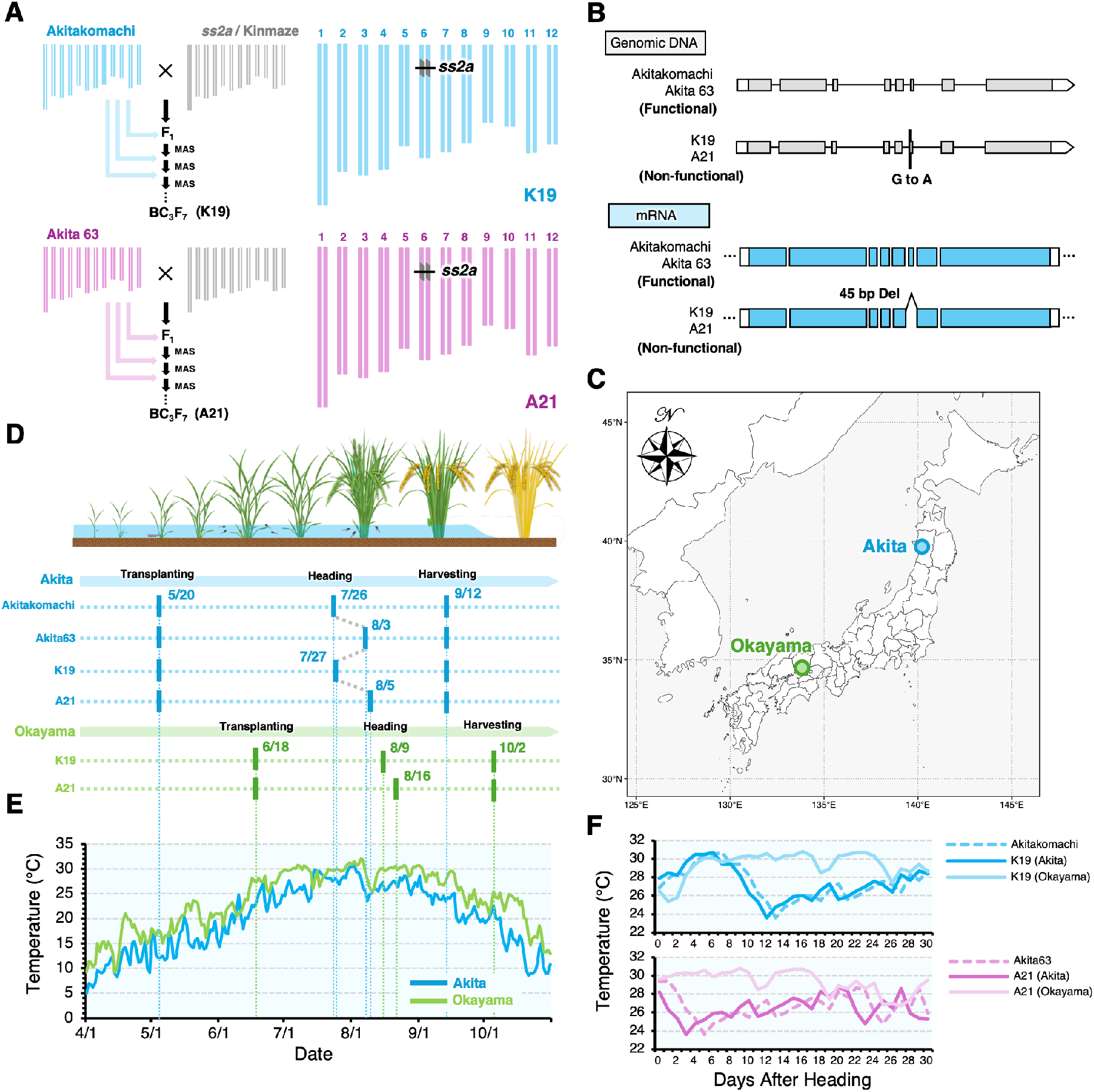
Experimental design, growth locations and environmental conditions used for evaluating the *ss2a* backcrossed inbred lines (BILs). (A) Development of the *ss2a* backcrossed inbred lines (BILs) in the genetic backgrounds of ‘Akitakomachi’ and ‘Akita 63’. Marker-assisted selection (MAS) was used during three successive backcross generations to introduce the non-functional *ss2a* allele, resulting in the BC_3_F_7_ lines K19 (‘Akitakomachi’ background) and A21 (‘Akita63’ background). Graphical genotypes of the selected BILs are shown on the right. Colored regions indicate the recurrent parent genome, whereas gray regions represent donor genome segments. The position of *ss2a* is indicated by the line. (B) Gene structure of *SSIIa* in the functional (‘Akitakomachi’ and ‘Akita 63’) and non-functional (K19 and A21) alleles. Boxes and lines indicate exons and introns, respectively. (C) Locations of the experimental fields in Akita (Katagami city) and Okayama (Kurashiki city), Japan. (D) Overview of the cultivation schedule at the two experimental sites. (E) Seasonal changes in daily mean air temperature during the growing season at the Akita (blue) and Okayama (green) experimental fields. Meteorological data were obtained from the AMeDAS stations in Akita City and Kurashiki City, respectively. (F) Daily mean air temperature during the grain-filling period from heading.

### Effects of growth location on polished-grain appearance and grain weight

Next, to determine the effects of the growth environment on grain traits, polished-grain appearance and brown-rice grain weight were evaluated for K19 and A21 grown at the two locations, Akita and Okayama. For the evaluation of polished-grain appearance, brown-rice grains were polished to approximately 90% of their original weight, and the proportion of grains exhibiting chalkiness was determined. (Fig. 2A–C). Grain chalkiness is promoted by environmental conditions such as high temperatures during grain filling and is one of the major factors responsible for deterioration of rice appearance quality. In addition, the *ss2a* mutation has also been reported to induce chalkiness of grains (Miura *et al*. 2018). In this study, polished grains harvested from each line at the two locations were classified into five categories according to the degree of chalkiness (Fig. 2C). Compared with their respective background cultivars, ‘Akitakomachi’ and ‘Akita 63’, both K19 and A21 showed reduced proportions of non-chalky grains (Fig. 2C). When the two growth locations were compared, K19 grown in Okayama showed a lower proportion of chalky grains than K19 grown in Akita. In contrast, A21 tended to show a lower proportion of chalky grains when grown in Akita than when grown in Okayama. In addition to appearance quality, brown-rice grain weight was measured to evaluate the degree of grain filling. In K19, no significant difference in grain weight was observed between seeds produced in Akita and Okayama, indicating relatively stable grain weight across the two growth locations (Fig. 2D). However, K19 grown in both Akita and Okayama showed significantly lower grain weight than its wild-type background cultivar, ‘Akitakomachi’ (Fig. 2D). In contrast, the grain weight of A21 grown in Akita was significantly higher than that of A21 grown in Okayama (Fig. 2D). When compared with its wild-type background cultivar, ‘Akita 63’, A21 grown in Akita showed a comparable grain weight, whereas A21 grown in Okayama exhibited significantly lower value (Fig. 2D).

**Fig 2.**
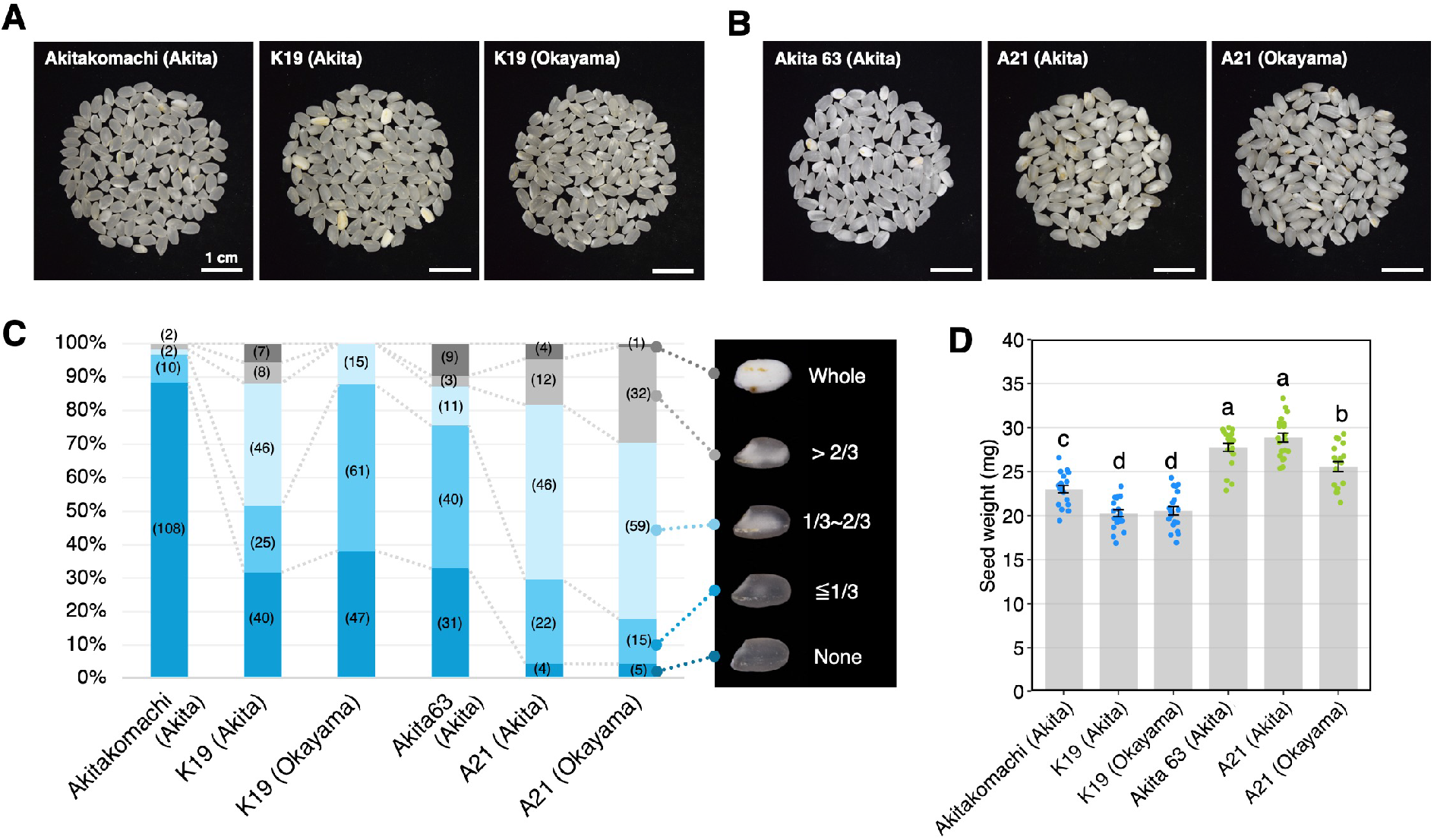
Seed appearance and seed weight of *ss2a* mutant lines grown in Akita and Okayama. (A, B) Representative images of polished rice grains of K19 (A) and A21 (B) grown in Akita and Okayama, together with their respective parental cultivars, ‘Akitakomachi’ and ‘Akita 63’ grown in Akita. (C) Distribution of chalkiness severity in polished rice grains. Grains were classified into five categories according to the proportion of the chalky area: None, ≤1/3, 1/3–2/3, >2/3, and Whole. Numbers in bars represent the number of seeds. (D) Seed weight of each line. Each dot represents an individual seed, and gray bars indicate the mean. Error bars indicate standard errors. Different letters indicate significant differences among groups according to Tukey’s multiple comparison test (*P* < 0.05).

### The ss2a mutation maintained a high proportion of short amylopectin chains under high-temperature conditions

Next, to determine the effects of the *ss2a* mutation on starch structure and its response to growth under high-temperature conditions, amylopectin chain-length distributions were analyzed by capillary electrophoresis. Compared with their respective wild types, both K19 and A21 showed increased proportions of short amylopectin chains (*i*.*e*. DP 5–10, with a peak at DP 8) at both locations (Fig. 3A, B). These alterations in chain-length distribution were consistent with the previously reported characteristics of SSIIa deficiency, in which impaired elongation of intermediate-length chains results in an increased proportion of short amylopectin chains (Miura *et al*. 2018). Notably, the proportion of chains at DP 8 was higher in Akita than in Okayama of both lines. When the effects of growth location were examined, K19 grown in Akita showed an increased proportion of short amylopectin chains with DP ≤ 10 and a decreased proportion of intermediate-to-long chains with DP > 10 compared with K19 grown in Okayama (Fig. 3C). A21 grown in Akita showed an increased proportion of short amylopectin chains with DP ≤ 15 and a decreased proportion of intermediate-to-long chains with DP > 15 compared with A21 grown in Okayama (Fig. 3C). As described above, although the changes in DP10-15 differed slightly between K19 and A21, an increase in short chains and a decrease in medium-to-long-chains were common to both genetic backgrounds. Kato *et al*. (2019) analyzed amylopectin chain-length distributions in ‘Akitakomachi’ grown under low- and high-temperature conditions (Fig. 3D). Their results showed that low-temperature conditions increased the proportions of amylopectin chains with DP ≤ 15, except for DP 8, while decreasing the proportions of intermediate-to-long chains with DP ≥ 16 (Fig. 3D). Since the changes observed in K19 and A21 between Akita and Okayama are generally similar to these changes, this strongly suggests that the endosperm starch of rice grown in Okayama is affected by high-temperature ripening.

**Fig 3.**
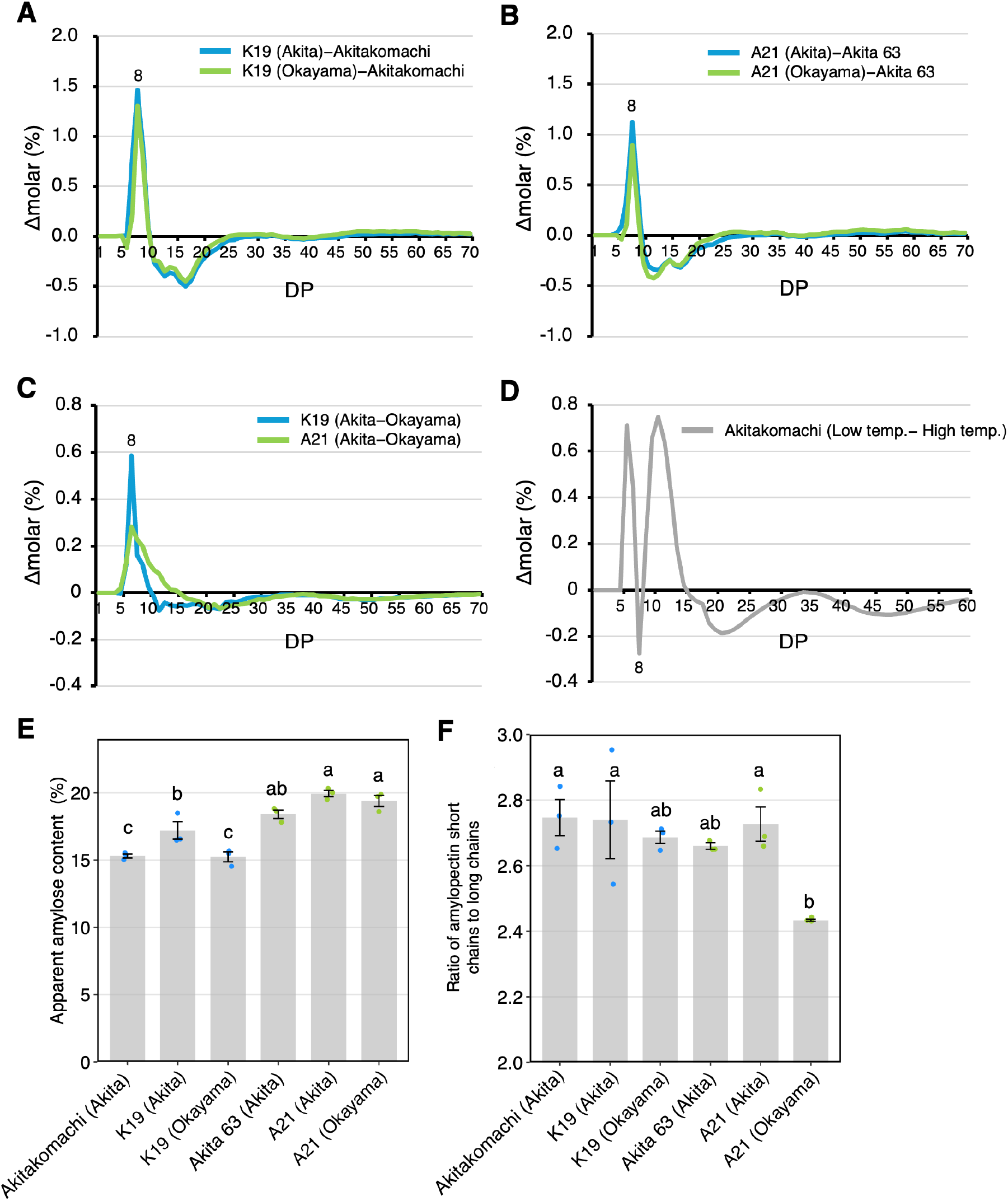
Starch structure of *ss 2 a* mutant lines grown in Akita and Okayama. (A–D) Differences in amylopectin chain-length distribution. (A) Differences between K19 grown in Akita or Okayama and ‘Akitakomachi’ grown in Akita. (B) Differences between A21 grown in Akita or Okayama and ‘Akita 63’ grown in Akita. (C) Differences between Akita- and Okayama-grown K19 and A21. The data represent the mean of three replicates, with standard error of less than 4% for all values. (D) Difference in the amylopectin chain-length distribution of ‘Akitakomachi’ grown under low- and high-temperature conditions. Data in (D) were adapted from Kato *et al*. (2019). Differences in chain-length distribution are expressed as Δmolar (%) for each degree of polymerization (DP). (E) Apparent amylose content of each line grown in Akita and Okayama. (F) Ratio of short to long amylopectin chains. Bars represent the mean, dots represent individual measurements, and error bars indicate standard errors. Different letters indicate significant differences among groups according to Tukey’s multiple comparison test (*P* < 0.05). Data in (E) and (F) were derived from the gel-filtration chromatography of debranched starch (Supplementary Fig. S1).

Analysis of starch gelatinization properties revealed that both K19 and A21 exhibited significantly lower onset temperature, peak temperature, and conclusion temperature than their respective background cultivars (Table 1). With regard to gelatinization enthalpy, “Akitakomachi” and “Akita 63” tended to have higher values than K19 and A21, respectively, but no significant difference was observed between the two growth locations (Table 1). Furthermore, within both K19 and A21, samples produced in Akita showed lower gelatinization temperatures than those produced in Okayama. In general, a higher proportion of short amylopectin chains with DP < 13 is associated with a lower starch gelatinization temperature (Hayashi *et al*. 2015, Hizukuri 1986). Therefore, these gelatinization properties were consistent with the results of the amylopectin chain-length distribution analysis. Collectively, these results suggest that although high temperatures during grain filling decrease the proportion of short amylopectin chains, *ss2a* mutant lines retain a higher proportion of short amylopectin chains than their respective wild types.

**Table 1.** Thermal properties of endosperm starch as determined by differential scanning calorimetry.

| Sample | T<sub>0</sub> (°C) | T<sub>p</sub> (°C) | T<sub>c</sub> (°C) | ΔH (mJ/mg) |
| --- | --- | --- | --- | --- |
| Akitakomachi (Akita) | 59.8±0.1<sup>a</sup> | 66.1±0.0<sup>a</sup> | 72.8±0.0<sup>c</sup> | 13.9±0.6<sup>a</sup> |
| K19 (Akita) | 52.9±0.1<sup>d</sup> | 61.4±0.1<sup>d</sup> | 69.6±0.1<sup>d</sup> | 13.0±0.2<sup>a</sup> |
| K19 (Okayama) | 57.0±0.1<sup>b</sup> | 64.9±0.1<sup>b</sup> | 73.6±0.2<sup>b</sup> | 13.4±0.4<sup>a</sup> |
| Akita 63 (Akita) | 60.2±0.1<sup>a</sup> | 66.0±0.0<sup>a</sup> | 73.1±0.1<sup>bc</sup> | 14.7±0.5<sup>a</sup> |
| A21 (Akita) | 54.4±0.0<sup>c</sup> | 61.8±0.1<sup>d</sup> | 70.0±0.0<sup>d</sup> | 12.8±0.7<sup>a</sup> |
| A21 (Okayama) | 56.8±0.1<sup>b</sup> | 64.3±0.0<sup>c</sup> | 74.5±0.2<sup>ab</sup> | 13.7±0.2<sup>a</sup> |

Mean value ± SE (n=3) of three seeds. T<sub>0</sub>, Onset temperature; T<sub>p</sub>, peak temperature; T<sub>c</sub>, conclusion temperature; ΔH, gelatinization enthalpy of starch. Different letters indicate significant differences among groups according to Tukey’s multiple comparison test ( $P < 0.05$ )

High temperatures during grain filling are known to reduce the expression of *GBSSI*, resulting in decreased amylose content(Larkin and Park 1999). Therefore, the apparent amylose content of the materials used in this study was compared by gel-filtration chromatography (Fig. 3E). In K19, samples produced in Okayama showed lower apparent amylose content than those produced in Akita, and K19 grown in Akita showed a higher apparent amylose content than ‘Akitakomachi’ grown in Akita (Fig. 3E). In contrast, A21 showed comparable apparent amylose contents between Akita and Okayama, both of which tended to be slightly higher than that of the background cultivar ‘Akita 63’ (Fig. 3E).

These results suggest that the *ss2a* mutation may compensate for the reduction in amylose content caused by high-temperature grain filling. Gel-filtration chromatography of debranched starch showed that the ratio of short to long amylopectin chains was significantly decreased in A21 grown in Okayama, while K19 showed a similar decreasing trend (Fig. 3F). In contrast, no significant differences were observed between the *ss2a* mutant lines and their respective WT backgrounds. These results suggest that the *ss2a* mutation may compensate for the reduction in amylose content caused by high-temperature grain filling.

### Sensory evaluation of cooked rice from the ss2a mutant lines

Sensory evaluations of cooked rice from K19 grown in Akita and Okayama were conducted since K19 is derived from ‘Akitakomachi’, a high-eating-quality rice cultivar in Japan (Fig. 4). Six sensory attributes and overall eating quality were evaluated for both freshly cooked and cooled rice. For each serving condition, ‘Akitakomachi’ grown in Akita was used as the reference and assigned a score of 3. K19 grown in Akita received higher sensory scores than K19 grown in Okayama under both serving conditions (Fig. 4).

**Fig 4.**
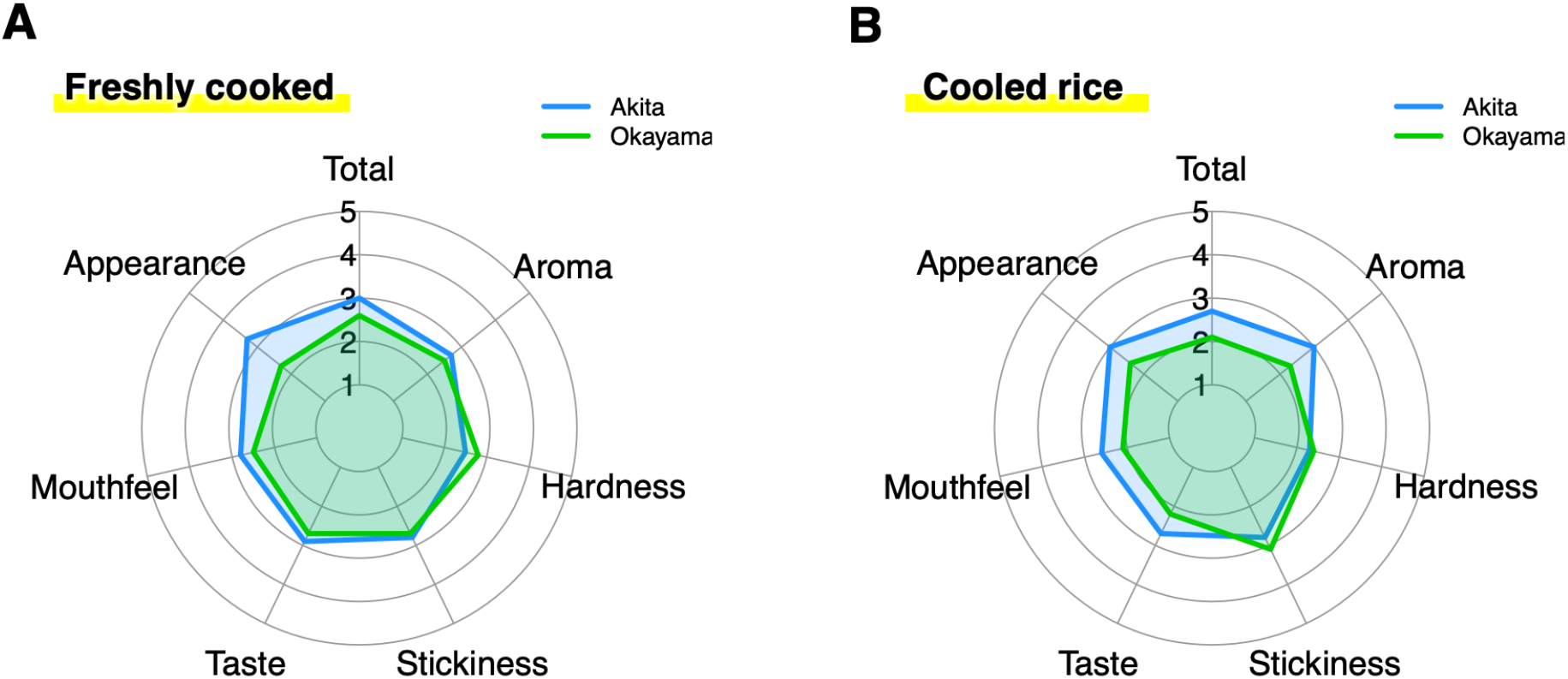
Sensory evaluation of K19 cooked rice from plants grown under different growth locations. Sensory attributes of K19 cooked rice were evaluated based on six parameters: appearance, aroma, hardness, mouthfeel, stickiness, and taste. Blue and green areas represent K19 grown in Akita and Okayama, respectively. Higher scores indicate higher ratings for each sensory attribute; for hardness and stickiness, higher scores indicate softer and stickier rice, respectively. The commonly consumed non-glutinous rice cultivar ‘Akitakomachi’ grown in Akita was used as a control, and its score was set at three.

## Discussion

High temperature during grain filling is one of the major environmental factors threatening the stability of rice yield and quality. Previous studies on the deterioration of rice quality caused by high-temperature grain filling have extensively investigated its effects on grain appearance, particularly the increased occurrence of chalky grains. However, high temperature affects not only the amount of starch accumulated in the endosperm but also its molecular structure. In particular, high-temperature conditions are known to decrease amylose content and the proportion of short amylopectin chains while increasing the proportion of intermediate-to-long amylopectin chains(Asaoka *et al*. 1984, Kato *et al*. 2019). Because the chain-length distribution of amylopectin is an important determinant of starch gelatinization properties and the retrogradation of cooked rice, such changes in starch molecular structure are likely to contribute to the deterioration of eating quality under high-temperature grain-filling conditions. Importantly, changes in starch molecular structure do not necessarily coincide with the formation of chalky grains. Therefore, suppressing chalkiness alone may not be sufficient to fully prevent the deterioration of rice quality under high-temperature conditions. In addition to improving grain appearance, breeding strategies that maintain an appropriate starch structure under high-temperature conditions are required.

To address this issue, SSIIa, which regulates the chain-length distribution of amylopectin, was chosen as a target. SSIIa is one of the major starch synthases responsible for elongating relatively short amylopectin chains during starch biosynthesis. Reduced SSIIa function suppresses the elongation of short chains, resulting in an increased proportion of short amylopectin chains, particularly those with DP 6–12 (Miura *et al*. 2018). This alteration occurs in the opposite direction to the decrease in short amylopectin chains associated with reduced BEIIb expression during high-temperature grain filling. Therefore, it was hypothesized that shifting the amylopectin chain-length distribution toward shorter chains in advance through the *ss2a* mutation would enable a relatively high proportion of short chains, even under high-temperature conditions. In other words, rather than completely preventing environmentally induced changes in starch structure, our strategy was to genetically shift the initial starch structure in the opposite direction so that the final starch structure remains within a desirable range under high-temperature conditions.

To test this hypothesis, two *ss2a* mutant lines, K19 and A21, with different genetic backgrounds were grown at two locations, Akita and Okayama, which differed in temperature during the grain-filling period (Fig. 1). From August to early September in 2024, corresponding approximately to the grain-filling period, average temperatures in Okayama were 2–4°C higher than those in Akita, allowing us to evaluate the effects of contrasting grain-filling temperatures under field conditions. Amylopectin chain-length distribution analysis showed that both K19 and A21 had higher proportions of short chains with DP ≤ 10 than their respective background cultivars grown in Akita (Fig. 3). This result was consistent with the known effect of the *ss2a* mutation(Miura *et al*. 2018) and, importantly, the increase in short chains was maintained even when the plants were grown in the warmer environment of Okayama. Thus, the effect of the *ss2a* mutation on amylopectin chain-length distribution was not abolished by the grain-filling environment, demonstrating that this mutation can genetically maintain the chain-length distribution toward shorter chains even under high-temperature conditions.

**Supplementary Fig. S1.**
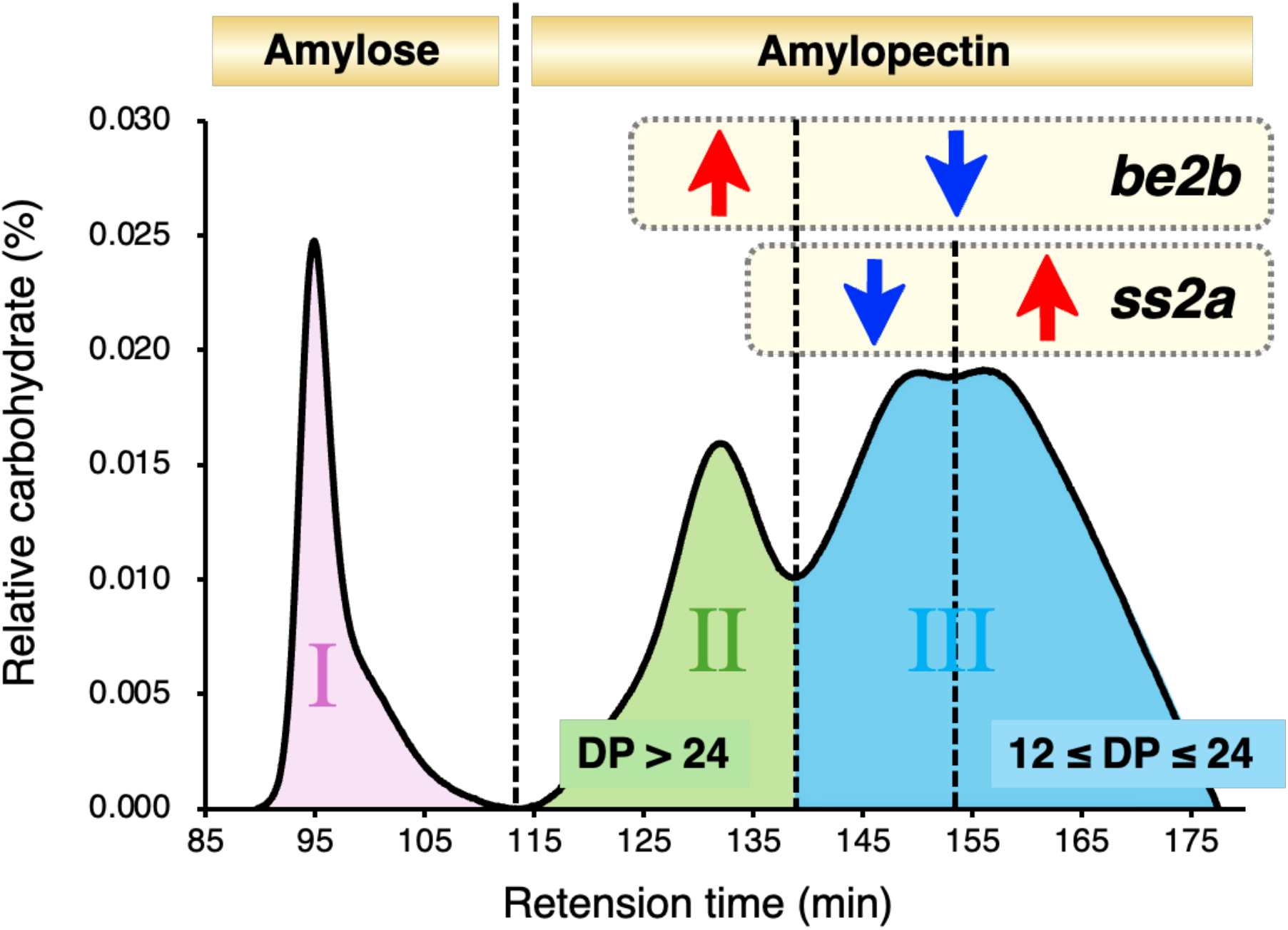
Schematic representation of starch fractionation by gel-filtration chromatography. A representative gel-filtration chromatogram of debranched starch is shown. The three major fractions were defined as Fractions (Fr.) I, II, and III. Fr. I mainly represents amylose, whereas Fr. II and Fr. III represent amylopectin chains with degrees of polymerization (DP) of 12–24 and >24, respectively. Reduced or deficient BEIIb activity increases Fr. II and decreases Fr. III, resulting in an increased ratio of Fr. II to Fr. III (II/III), which represents the relative abundance of short to long amylopectin chains. In contrast, reduced or deficient SSIIa activity decreases the earlier-eluting portion and increases the later-eluting portion within Fr. III. Because these opposing changes occur within the same fraction, they largely offset each other and therefore have little effect on the II/III ratio.

When the present results were compared with those reported by Kato *et al*. 2019, the overall pattern of differences between the low- and high-temperature (Akita and Okayama, respectively in the Fig. 3C) conditions were largely conserved (Fig. 3C and 3D). However, an interesting difference was observed in the short-chain region. In the *ss2a* mutant lines, the short-chain region showed a single peak centered at DP 8, whereas the previous report using WT exhibited a bimodal pattern, with a pronounced trough at DP 8 separating the two peaks (Fig. 3C and 3D). This contrasting response at DP 8 may be explained by the different roles of SSI and SSIIa in amylopectin chain elongation. SSI primarily elongates short glucan chains of DP 6–7 to DP 8–12(Fujita *et al*. 2006), and *SSI* expression has been reported to increase under high-temperature conditions (Kato *et al*. 2019, Yamakawa *et al*. 2007). Thus, enhanced SSI activity under high-temperature conditions could explain the particularly pronounced decrease at DP 8 in the low-versus high-temperature difference observed in the WT (Fig. 3D). In contrast, SSIIa normally elongates short chains into intermediate-length chains. Thus, in the *ss2a* mutants, the absence of functional SSIIa may have caused the effect of high temperature to be particularly pronounced around DP 8, resulting in the characteristic response observed in the mutants (Fig. 3C). For longer chains (DP ≥ 15), the response to temperature was similar between the *ss2a* mutants and WT. This similarity may reflect a common response to high temperature, particularly the reported reduction in *BEIIb* expression, which would affect amylopectin branching and consequently increase the relative abundance of longer chains. Thus, the *ss2a* mutation does not completely eliminate the temperature-dependent alteration in amylopectin chain-length distribution. Nevertheless, the mutant lines grown in Okayama still maintained higher proportions of short chains than their respective background cultivars. The *ss2a* mutation may therefore shift the final chain-length distribution toward shorter chains even after the proportion of short chains has been reduced by high-temperature grain filling. This finding supports our proposed strategy of compensating for an environmentally induced change in starch structure through a genetic alteration that acts in the opposite direction.

These alterations in amylopectin chain-length distribution were also clearly reflected in the thermal gelatinization properties of starch. Compared with their respective background cultivars, K19 and A21 exhibited lower onset, peak, and conclusion temperatures of gelatinization (Table 1). Notably, the gelatinization temperatures of K19 and A21 grown in Okayama were lower than those of ‘Akitakomachi’ and ‘Akita 63’ grown in Akita, respectively. These results suggest that the introduction of the *ss2a* mutation more than compensated for the negative effects of high-temperature grain filling on starch properties. Amylopectin side chains form double helices with adjacent chains, and the assembly of these helices contributes to the crystalline regions within starch granules. In general, an increased abundance of short amylopectin chains reduces the thermal stability of these double helices and crystalline structures, thereby promoting starch gelatinization at lower temperatures(Hayashi *et al*. 2015, Hizukuri 1986). Thus, the increased proportion of short chains in the *ss2a* mutant lines is consistent with their lower gelatinization temperatures. Furthermore, within each mutant line, starch from plants grown in Akita showed lower gelatinization temperatures than starch from plants grown in Okayama, consistent with the higher proportion of short amylopectin chains in the Akita-grown samples (Table 1 and Fig. 3). These results indicate that differences in amylopectin chain-length distribution caused by the *ss2a* mutation and by the grain-filling environments were directly reflected in the thermal gelatinization properties of starch.

Apparent amylose content was also affected by both the *ss2a* mutation and the grain-filling environment. High-temperature grain filling is generally known to reduce GBSSI expression, resulting in decreased amylose content in japonica cultivars (Larkin and Park 1999). Consistent with this response, the apparent amylose content of K19 grown under the warmer conditions in Okayama was lower than that of K19 grown in Akita (Fig. 3E). On the other hand, the *ss2a* mutation has been reported to increase apparent amylose content (Miura et al., 2018), and in the present study the mutant lines showed apparent amylose contents that were higher than, or at least comparable to, those of their respective background cultivars (Fig. 3E). In particular, A21 showed no clear reduction in apparent amylose content between Akita and Okayama, and the value remained slightly higher than that of its background cultivar even in Okayama. Comparison of mean temperature profiles relative to the flowering date of each line showed that the temperature difference between Akita and Okayama was greater during the later stage of grain filling in K19, whereas it was greater during the earlier stage in A21 (Fig. 1F). This difference in the timing of exposure to higher temperatures may partly explain the different responses of apparent amylose content between K19 and A21(Fig. 3E). In particular, the reduction in apparent amylose content observed in K19, but not clearly in A21, may suggest that exposure to high temperatures during the later stage of grain filling is particularly important for determining the final amylose content. This possibility is consistent with the finding of Asai *et al*. (2014) that amylose accumulation continues to increase after 20 days after flowering until grain maturity under field condition(Asai *et al*. 2014). These observations raise the possibility that the *ss2a* mutation may partially compensate not only for high-temperature-induced changes in amylopectin chain-length distribution but also for the reduction in apparent amylose content.

Gel-filtration chromatography generally separates debranched starch into three major fractions, designated Fractions (Fr.) I, II, and III (Supplementary Fig. S1). Fr. I mainly represents amylose, whereas Fr. II and Fr. III represent amylopectin chains with degrees of DP of 12–24 and >24, respectively. The Fr. II/III ratio can therefore be used as an indicator of the relative abundance of short to long amylopectin chains. In the present study, our results showed that the Fr. II/III ratio was significantly decreased in A21 grown in Okayama, while K19 showed a similar decreasing trend (Fig. 3F). This response may reflect reduced BEIIb expression in high temperature and was consistent with the results of the amylopectin chain-length distribution analysis (Fig. 3E). Reduced or deficient BEIIb activity increases Fr. II and decreases Fr. III, resulting in an increased Fr. II/Fr. III ratio, which represents the relative abundance of short to long amylopectin chains (Supplementary Fig. S1). In contrast, no significant differences in the Fr. II/III ratio were observed between the *ss2a* mutant lines and their respective WT backgrounds (Fig. 3F). SSIIa primarily elongates short amylopectin chains of DP ≤12 to intermediate-length chains of DP 13–24. Because the changes caused by reduced SSIIa activity occur mainly within the same gel-filtration fraction (Fr. III), the *ss2a* mutation may alter the chain-length distribution within this fraction without substantially affecting the Fr. II/Fr. III ratio (Fig. 3F and Supplementary Fig. S1).

The effects of the *ss2a* mutation on grain appearance were less straightforward than those on starch molecular structure (Fig. 2). Both K19 and A21 showed lower proportions of perfect grains than their respective background cultivars, consistent with the previous observation that the *ss2a* mutation itself can induce grain chalkiness (Miura *et al*. 2018) (Fig. 2A–C). Moreover, the effect of growth location differed depending on the genetic background: K19 showed less chalkiness when grown in Okayama, whereas A21 tended to show less chalkiness when grown in Akita (Fig. 2A–C). Similarly, grain weight did not differ significantly between locations in K19, whereas A21 produced heavier grains in Akita than in Okayama (Fig. 2D). These results suggest that chalkiness and grain filling are not determined solely by grain-filling temperature but are influenced by genotype-by-environment interactions as well as differences in developmental characteristics, such as heading date and the duration of grain filling. In particular, because the present study was conducted as a field experiment at two geographically distinct locations rather than under temperature-controlled conditions, environmental factors other than temperature, including solar radiation, diurnal temperature variation, and soil and water conditions, may also have differed between the sites. In the present study, K19 headed approximately one week earlier than A21 (Fig. 1D). Therefore, the grain-filling periods of the two lines did not completely overlap, and the temperature conditions experienced during grain filling likely differed between them. Previous study using *ss2a* rice lines has demonstrated that differences in heading date alter the temperature conditions experienced during grain filling and consequently affect grain characteristics, including starch properties and grain yield(Crofts *et al*. 2022). Thus, when interpreting the phenotypic differences observed between K19 and A21, it is necessary to consider not only differences in their genetic backgrounds but also the potential effects of differences in grain-filling temperatures resulting from their different heading dates.

The sensory evaluation of K19 revealed differences in eating quality between the two cultivation locations (Fig. 4). Rice produced in Akita received higher sensory scores than rice produced in Okayama for both freshly cooked and cooled rice. This tendency corresponded with the starch characteristics of the Akita-grown samples, which had higher proportions of short amylopectin chains and higher apparent amylose content (Fig. 3). Because starch with a lower gelatinization temperature (Table 1) tends to retrograde more slowly after cooking, this characteristic may have contributed to maintaining the eating quality of K19 after cooling. These results suggest that the altered starch structure associated with the *ss2a* mutation may influence not only the structure and physicochemical properties of starch but also the eating quality of cooked rice, particularly after cooling. Based on the starch structural and thermal gelatinization properties, the *ss2a* mutation appeared to compensate for the negative effects of high-temperature grain filling (Fig. 3C, E, F; Table 1). Therefore, we expected K19 grown in Okayama to show eating quality comparable to that of the control cultivar ‘Akitakomachi’. However, the sensory evaluation did not support this expectation, as K19 grown in Okayama received lower scores than ‘Akitakomachi’ (Fig. 4). This discrepancy indicates that the eating quality of cooked rice cannot be explained solely by starch structure and gelatinization properties. Rather, it is determined by multiple factors, including grain size, polishing degree associated with grain size, amylose content, and the amount of water added during cooking.

Overall, our results suggest a new breeding strategy for adaptation to high-temperature grain filling: genetically shifting starch structure toward a higher proportion of short amylopectin chains to counteract the changes induced by heat stress. Conventional breeding for tolerance to high-temperature grain filling has primarily focused on suppressing the formation of chalky grains or maintaining starch accumulation under high-temperature conditions. In contrast, our results indicate that shifting amylopectin chain-length distribution toward shorter chains through the *ss2a* mutation may allow a relatively high proportion of short chains to be retained even after their abundance is reduced by high-temperature grain filling. However, the results of this study are based on a single year of field experiments. Further studies over multiple years are therefore needed to determine whether similar effects can be reproduced under different climatic conditions. In addition, the effects of the *ss2a* mutation on agronomic traits, including yield and yield related components, should be evaluated for its practical use in rice breeding. Such evaluations should be conducted together with analyses of starch structure and eating quality. Furthermore, SSIIa elongates amylopectin chains via SSI elongation using the non-reducing ends of branches generated by BEIIb as substrates(Abe *et al*. 2014, Crofts *et al*. 2017). Therefore, reduced *BEIIb* expression is a more direct cause of the changes in amylopectin chain-length distribution under high-temperature conditions. Thus, identifying *BEIIb* alleles that maintain stable expression under high-temperature conditions may provide a more fundamental solution to this problem. Such alleles will represent valuable genetic resources for breeding rice with stable starch properties under high-temperature conditions.

## Author Contribution Statement

NF conceived and designed the study. NF, SH, SM, NC performed the experiments and conducted formal analyses. YH and NO provided technical assistance to NF. RM and TY were responsible for plant cultivation and data collection in Okayama. SH prepared the figures and table, and drafted the manuscript. NF and SH revised and finalized the manuscript. All authors have read and approved the final version of the manuscript.

## Acknowledgements

The authors thank Ms. Yuko Nakaizumi (Akita Prefectural University) for growing the rice plants, and Nagase Viita Co., Ltd. for providing isoamylase.

